# Mapping Single-molecule Protein Complexes in 3D with DNA Nanoswitch Calipers

**DOI:** 10.1101/2023.07.10.548386

**Authors:** Prakash Shrestha, Darren Yang, William M. Shih, Wesley P. Wong

**Affiliations:** Program in Cellular and Molecular Medicine, Boston Children’s Hospital; Wyss Institute for Biologically Inspired Engineering, Harvard University; Department of Cancer Biology, Dana-Farber Cancer Institute; Department of Biological Chemistry and Molecular Pharmacology, Blavatnik Institute, Harvard Medical School

## Abstract

The ability to accurately map the 3D geometry of single-molecule complexes in trace samples would lead to new insights into molecular mechanics and provide an approach for single-molecule structural proteomics. To enable this, we have developed a high-resolution force-spectroscopy method capable of measuring multiple distances between labeled sites in natively folded protein complexes. Our approach combines reconfigurable nanoscale devices we call DNA Nanoswitch Calipers, which we have previously introduced, with a force-based barcoding system to distinguish each measurement location. We demonstrate our approach by reconstructing the tetrahedral geometry of biotin-binding sites in natively folded streptavidin, with 1.5-2.5 *Å* agreement to previously reported structures.

## Main

The ability to determine the spatial organization of residues in biomacromolecular complexes has led to a deeper understanding of biological mechanisms and rational approaches for developing therapeutics^1,2^. Despite recent developments in computational tools for predicting protein structure^3–5^, challenges remain in their ability to predict the assembly of multi-protein complexes, mechanical properties and conformational transitions. Experimentally, technologies such as super-resolution microscopy^6–9^ and structural biology methods such as cryoEM^10–12^ have opened new avenues for molecular measurement. Yet application of these powerful techniques in fields such as proteomics has been constrained due to limitations in resolution, accuracy, and throughput on one hand, and the challenge of working with trace, heterogeneous samples on the other. Emerging single-molecule approaches have the potential to circumvent these hurdles. Single-molecule Fluorescence Resonance Energy Transfer (smFRET)-based systems^13–16^ as well as single-molecule force spectroscopy^17–20^ and nanopore^21^ approaches have been used to study the shape of biomolecules and the dynamics of their structural transitions. For example, notable AFM studies have demonstrated the reconstruction of folded protein geometry^22,23^, although these measurements were largely limited to a single pairwise distance per molecule. Resolving the challenge of making multiple atomically precise distance measurements within natively-folded single biomolecular complexes could enable new applications in single-molecule proteomics, including the determination of three-dimensional geometry from trace biological samples and enable studies of molecular interactions, heterogeneity, and structure-function relationships.

To enable this measurement capability, we have developed an approach that combines reconfigurable nanodevices with single-molecule force spectroscopy. Previously, we introduced the concept of DNA Nanoswitch Calipers (DNC), and demonstrated its application to protein-sequence fingerprinting by measuring the distances between specific residues within denatured peptides that have been chemically labeled with DNA hanldes^24^. As described in Fig. 1, each DNC consists of a DNA tether containing a pair of grabber sequences complementary to the DNA handles on an analyte of interest. When these grabbers hybridize to two handles on an analyte, a section of DNA loops out. When a sufficient force is applied to the DNC tether at its two ends, one grabber is released and the measured change in distance between the two ends of the tether under tension can be used to infer the original distance between the handles. In this present study, by combining DNC with force-based barcodes to augment our ability to disambiguate which labeled sites are being sampled, we measure the distances between multiple coordinates on the surface of an intact, natively folded protein complex, and reconstruct the three-dimensional geometry of these coordinates (Fig. 1). As a proof of concept, we reconstruct the geometry of biotin-binding sites in streptavidin by measuring the pairwise distances between binding sites and then using symmetry considerations to calculate their relative positions in 3D space. Streptavidin is a tetramer consisting of four identical, non-covalently attached units^25^ that exhibits a D2 symmetric structure (i.e. dimer of dimers), as shown by X-ray crystallography, in which the biotin-binding sites form an irregular tetrahedron with opposite sides that are equal in length^26,27^ (Fig. 2a).

**Figure 1.**
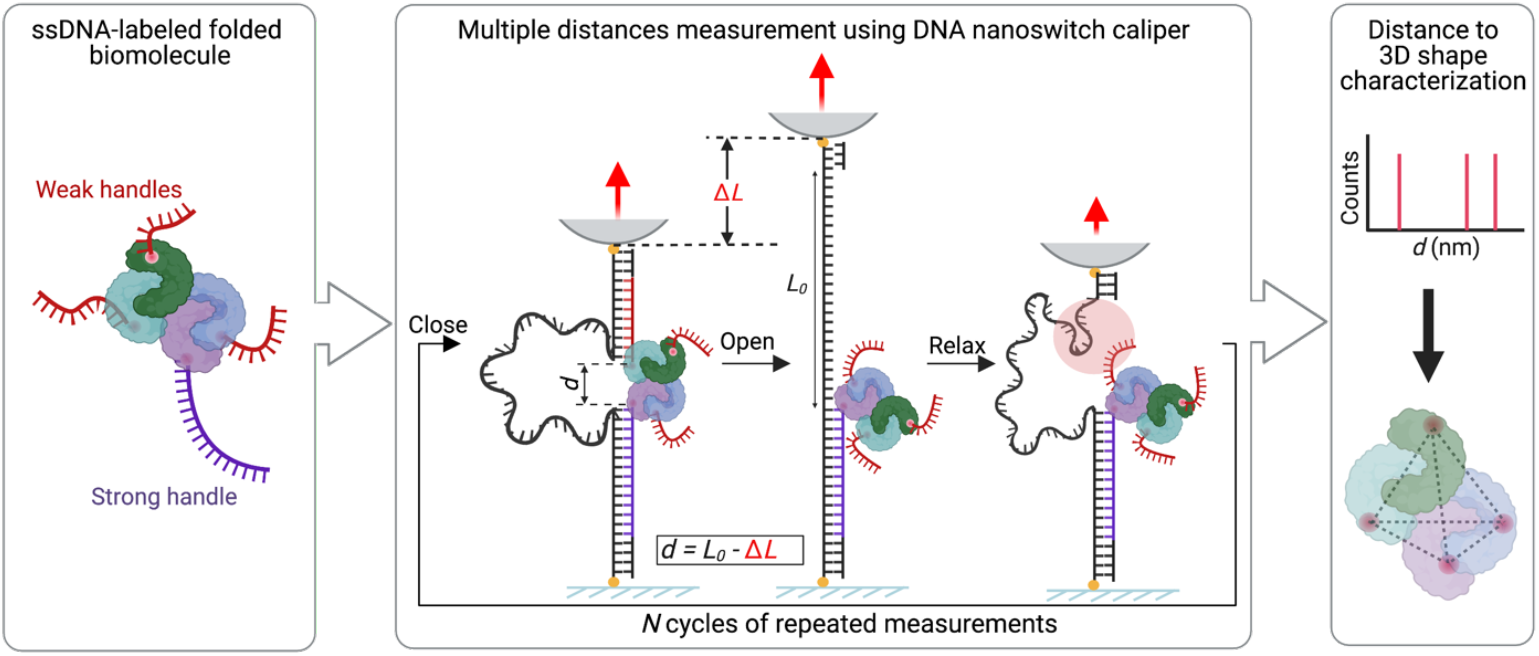
Schematic overview of single-molecule 3D mapping with DNA Nanoswitch Calipers (DNC) distance measurements. (*left*) The natively folded biomolecular complex is labeled with ssDNA handles, a strong DNA handle (purple) that serves as an anchoring site for DNC, and weaker ssDNA (red) handles. (*middle*) Multiple distances are measured between red and purple handles using DNC actuated by mechanical force. Each DNC has one DNA “grabber” complementary to the weak (red) handles, and one grabber complementary to the strong (purple) handle. Within each measurement cycle, the force is cycled between a low force and a high force. Under low force, the DNC forms a loop when both handles in the target are bound, and under high force the loop opens due to release of a weak handle. This process is repeated for N cycles of measurement with a red handle stochastically rebind each cycle at low force, closing the loop again. (*top right*) Distribution of the measured distances *d* calculated by subtracting the change-in-length *ΔL* from the loop length *L*_*0*_, which serves as a reference to convert changes in length to absolute distance. (*bottom right*) Reconstructed geometry for positions of DNA-labeled sites in the folded biomolecular complex.

**Figure 2.**
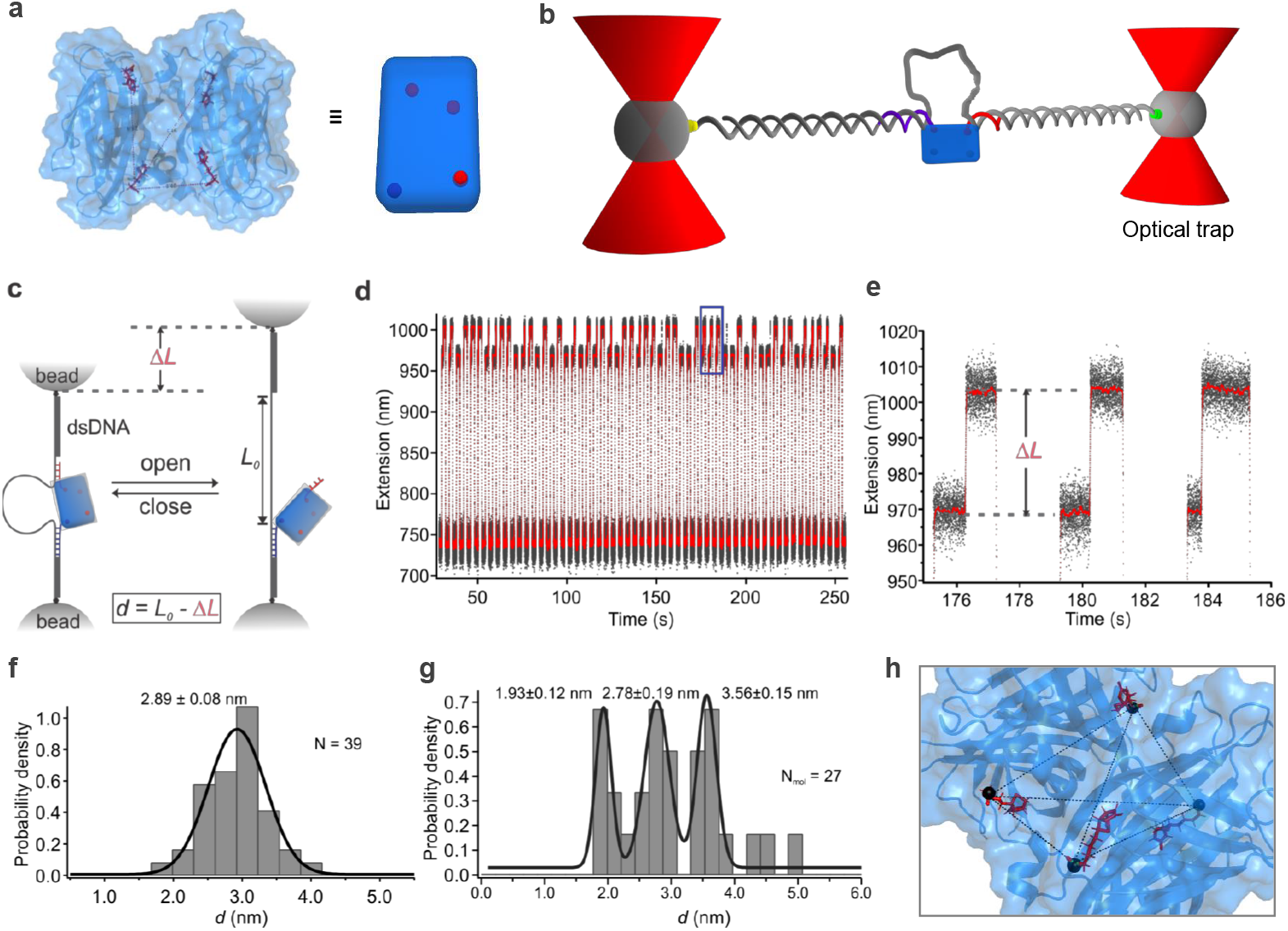
Single-molecule distance measurements with DNA Nanoswitch Calipers to determine the 3D geometry of binding sites in the streptavidin-biotin complex. **a**, Structure of streptavidin-biotin complex (PDB 6M9B) (*left*) with a schematic representation (*right*) where red spheres indicate biotins bound to streptavidin. **b**, Schematic representation of the DNC tethered between two optically trapped microbeads. The target streptavidin is bound to two biotinylated ssDNA handles each complementary to a region of the DNC, a longer handle (purple) provides strong attachment and shorter handle (red) that shears off the DNC at moderate force during measurement cycles. **c**, DNC measurement of the distance between two biotin-binding sites in a streptavidin molecule. The DNC is looped when the target streptavidin molecule is bound at both handles, becomes linear when the red handle shears of at a higher force, and reverts back to the looped configuration when the red handle rebinds at a lower force. **d**, Experimental graph showing changes in extension of the DNC over many cycles of low and high force (black curve indicates raw data, red curve indicates 200 points sliding window average). **e**, Zoomed in view of purple rectangle in **d**, showing change-in-extension due to unlooping of the DNC due to force. **f** Histogram of the absolute distances calculated by subtracting *ΔL* from the loop length (*L*_*0*_). **g** Histogram of the per-molecule average distances of 27 molecular complexes. **h**, Superposition of the DNC determined position of the carbon in the carboxylic group of each biotin (black spheres) on the crystal structure (PDB 6M9B).

As a first step to validating our approach, we used DNC to measure a single distance between pairs of biotin-binding sites for each single-molecule complex. First, we loaded one long biotinylated ssDNA strong handle and one short biotinylated ssDNA weak handle on each caliper and then mixed with tetrameric streptavidin for bivalent capture (Fig. 2 and Supplementary Fig. 1). The shearing force of the weak handle in our experiments (i.e., ∼26 pN) was significantly lower than the biotin-streptavidin unbinding force^28–31^, so that the biotin oligonucleotide generally remained attached to the streptavidin complex after each measurement cycle. Within each measurement cycle, force was applied to the ends of the DNC using optical tweezers and varied between a low force and a high force; at the high force of ∼26 pN, a sudden change in extension (*ΔL*) results from shearing of the weak handle, and at the low force of ∼0 pN, rebinding of the weak handle occurs, relooping the DNC tether. This process was repeated for several cycles as shown in figures 2 c-e.

The loop length (*L*_*0*_) was determined by performing calibration measurements of the DNC constructs at the distance measuring force^24^; the distance *d* was then calculated as *d = L*_*0*_ *– ΔL* (Fig. 2c). The distance distribution analyzed per molecule (Fig. 2f) obtained from several numbers of molecules clearly shows the three distances as expected (Fig. 2g and Supplementary Fig. 2). Using a prior assumption of irregular tetrahedron geometry from X-ray crystallography^32^, we used those distances to reconstruct the 3-dimensional geometry of the biotin binding sites of streptavidin, which closely matched the previously established structure, exhibiting an RMSD of 1.54 *Å* (Fig. 2h and Supplementary Fig. 3).

Next, to enable the measurement of multiple distances per single-molecule complex, we developed a barcoding system that enabled the identification of each binding site to be resolved from its unique force-extension behavior. This system increases the resolution of the DNA Nanoswitch Calipers by enabling sites that are closer than the native resolution of a single distance measurement to be distinguished and then resolved, with increased localization precision obtained by averaging repeated distance measurements, in the spirit of super-resolution approaches such as PALM^9^, FPALM^33^ and STORM^34^. Specifically, here we used hairpin barcodes in the weak shearing handles to help resolve all the distances between the biotin-binding sites in a single-streptavidin molecule. We incorporated a long, short, and “no hairpin” structure in the weak handle, in which the length due to unfolding of the hairpin serves as a barcode (Fig. 3 a,b). To functionalize streptavidin with biotinylated handles, we used a similar approach as discussed above but included a subsequent incubation with two other biotinylated weak shearing handles to have the streptavidin fully occupied (see Methods and Supplementary Fig. 4). We applied the following force ramping and jumping strategies in optical tweezers to measure the length of hairpin barcodes and the correspondingly measured distances of the biotin-binding sites in a streptavidin molecule. For example, we applied a slow ramping of the force up to ∼15 pN to probe the unfolding of the hairpin structure, and then jumped to a force (∼20 pN) to measure the distances of biotin-binding sites before jumping to a force (∼30 pN) for shear-opening the loop of DNC, and then jumped back to the distance measuring force (∼20 pN) (Fig. 3 c-e). The loop length (*L*_*0*_) of the DNC at the distance measuring force was determined as previously mentioned (Supplementary Fig. 5), and then used to calculate the distance *d*. By repetitive cycles of distance measurements in a single streptavidin-biotin complex, we observed the stochastic binding of all three differently barcoded weak-handles (Fig. 3e,f and Supplementary Fig. 6 and 7). As expected, we could identify in real time how many handles were bound per streptavidin and then precisely resolve all the distances of biotin-binding sites, which would otherwise be difficult.

**Figure 3.**
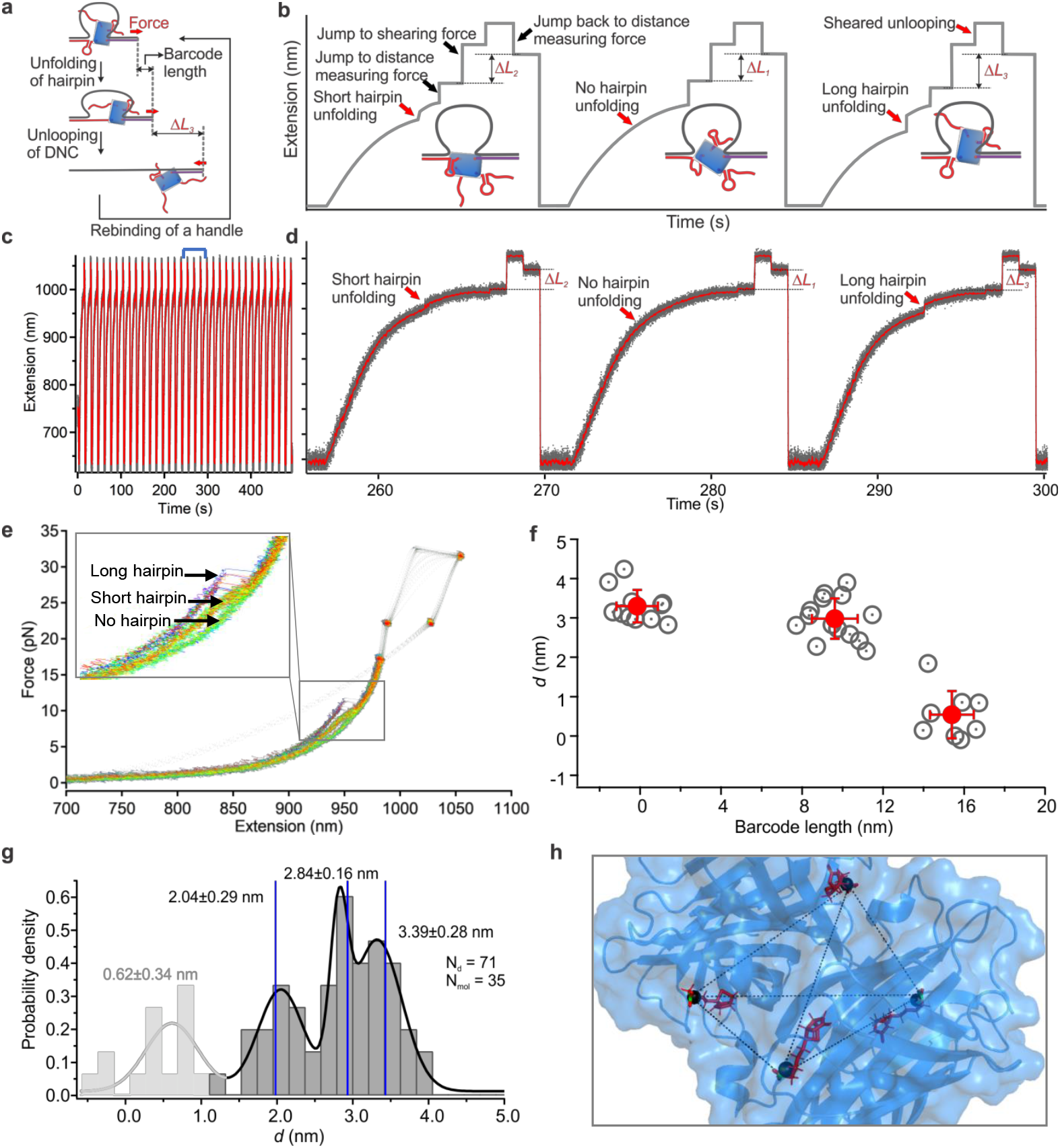
3D mapping of streptavidin-biotin binding sites using force-based barcodes to enable multiple DNC distance measurements on each molecular complex. **a**, Schematic showing measurement of multiple distances in a single streptavidin-biotin complex using hairpin barcodes incorporated into the weak shearing handles (shown in red). Initially, the DNC is stretched slowly until the hairpin opens, enabling handle identification from the change in length; next, the force is rapidly increased to initiate loop opening at a higher force, enabling the distance between handles to be measured as described in Figure 1; lastly, the force is dropped to enable rebinding to another handle. This process is repeated until all barcoded handles have been sampled, in order to measure all three distances in the streptavidin-biotin complex. **b** Outline of distance measurement strategy for optical tweezers. Force is ramped to ∼15 pN to detect the unfolding of hairpins, followed by a jump to a distance measuring force (∼20 pN) to measure the length of the tether, a jump to a higher force to shear off the weak handle and unloop the DNC, a jump back to the distance-measuring force (∼20 pN) to measure the change in length, followed by a jump to the rebinding force (∼0 pN). Three cycles are depicted to illustrate the unfolding signature of the three hairpin barcodes. **c**, Experimental graph of the extension vs. time for a DNC over multiple cycles of force application; gray and red curves indicate raw and smoothed data points, respectively. **d**, Zoom-in of a section (indicated by blue arrows) from (c) depicting the presence of all three barcodes in a row. **e**, Overlaid force vs. extension curves of repetitive cycles depicting the measurement of all three distances in a single streptavidin molecule. Gray arrows in the inset point to the unfolding transitions of the long, short and no hairpins in the weak handle. **f**, Scatter plot (gray circles with dots) of the measured length of the barcodes and the corresponding distances between the biotin-binding sites in a single molecule. The red-filled circles indicate the mean position of the corresponding populations and error bars indicate the standard deviation of the mean. **g**, Histogram depicting the aggregate of mean distances measured in several molecules. Blue vertical lines indicate the expected distances between biotin-binding sites based on the structure determined by X-ray crystallography (PDB 6M9B). **h**, Superposition of the determined position of carbon in the carboxylic group of each biotin (black spheres) on the crystal structure.

When we aggregated all the distances measured over different molecules that were bound with different numbers of DNA handles, the distance distribution showed three primary peaks at the expected distances (Fig. 3g). As in our previous example, by analyzing these three distances in light of the expected irregular tetrahedral symmetry, we were able to reconstruct the 3-dimensional geometry of the biotin-binding sites in streptavidin. And as before, the geometry of the binding sites determined by the DNC approach agreed closely with that determined by X-ray crystallography (RMSD 2.34 *Å*) (Fig. 3h and Supplementary Fig. 8).

Interestingly, however, we found that when a single streptavidin complex was fully occupied by all DNA handles, the distance between closely located biotin-binding sites appeared shorter than expected (Fig. 3f). To further investigate the existence of a shorter-than-expected distance in streptavidin complexes fully saturated by biotin, we performed additional experiments in which only one distance per complex was measured through attached DNA-biotin handles, while the remaining two biotin-binding sites were putatively occupied by biotin, through preincubation with a saturating concentration of DNA-free biotin in solution (Supplementary Fig. 9). Interestingly, we again found a population exhibiting shorter than expected distances (Supplementary Fig. 10 and 11), confirming our observation that when the streptavidin complex has all sites occupied it can exhibit subtly different physical properties than when only two sites are occupied. Comparing the aggregated single-distance results to the barcoded measurements, the three primary peaks in the distance distributions were similar and as predicted from X-ray crystallography, and both distributions had a smaller peak at a shorter distance, though this peak was slightly broader with a slightly higher peak value in the all-handle bound barcoded complex (Supplementary Fig. 11). Changes in the physical properties of fully-bound streptavidin are consistent with a previous study comparing the structures of free streptavidin and biotin-bound streptavidin, which showed that the helical loop near the respective biotin-binding sites had different configurations^27,35^; the loop has an open configuration and is relatively flexible in free streptavidin, whereas the loop has a closed configuration and is relatively compact when biotin is bound. Careful examination of the position of bound-biotin in the streptavidin complex shows that its carboxyl group remained slightly buried below the surface of the loop (Supplementary Fig. 12a). When considering the direction of force during caliper measurements and the positions of the biotin-attached linkers, we observe that when biotins attached to the caliper are located at the two longer distances, they are largely oriented along the pulling direction with the linkers directed away from the surface of streptavidin; however, when the biotins are positions at the shortest distance they are almost perpendicular to the pulling direction (Supplementary Fig. 12b) and the linkers must pull across the surface of streptavidin. This could lead to an apparent length of the linker that is shorter for the biotins bound to the closest binding sites, which would lead to a longer measured *ΔL*, and a shorter apparent *d*, depending on the flexibility of the protein. We speculate that when all binding sites are occupied, a decrease in the flexibility of the helical loop nearby the biotin-binding sites and an increase in compactness of the streptavidin complex may lead to an increase rigidity that exacerbates this effect.

In summary, we have developed a technique to measure multiple distances with barcodes to enhance the spatial resolution of DNC on a per-molecule basis. We measured the distances between biotin-binding sites in a streptavidin molecule using three hairpin barcodes as a proof-of-concept example. We successfully reconstructed the three-dimensional tetrahedron geometry of biotin-binding sites in the streptavidin molecule, which agrees closely (RMSD 2.34 *Å*) with the geometry determined from X-ray crystallography. DNC is a powerful and accessible technique to study the three-dimensional positions of residues in single biomolecules or biomolecular complexes that should lead to new applications in single-molecule proteomics, including the determination of three-dimensional molecular conformations from trace biological samples, and the discovery of new biomarkers for diagnostics and allosteric modulators for therapeutics.

## Methods

### Synthesis of DNA Nanoswitch Caliper (DNC) constructs

DNA nanoswitch calipers were synthesized as previously described^24^. Briefly, a long dsDNA handle was prepared by the PCR amplification of a 2820 bp region of the pET-26b (+) plasmid (see primer sequences in Supplementary table 1). After PCR and purification, the construct was digested with BsaI (New England Biolabs) to produce a sticky end to ligate with the other DNA fragment later. Next, the designed oligos such as short-loop oligonucleotide, 5′-digoxigenin modified oligonucleotide (DNC1), and other strands (DNC2 and DNC3) were mixed at equimolar concentration with the splint oligonucleotide at a 10-fold molar excess and annealed by slow ramping of temperature from 95 °C to 22 °C at the rate of -1°C/min. For DNC to measure multiple distances per molecule, oligonucleotides long-loop, DNC4 and DNC5 were added instead of short-loop, DNC1 and DNC2, respectively. The mixture was heated at 95 °C for 5 min and slowly annealed to room temperature. Then, the annealed mixture was ligated to the dsDNA construct prepared by PCR amplification using T4 DNA ligase (New England Biolabs) at 16°C for 16 h and the ligase was heat deactivated at 65 °C for 20 min. The fully-ligated products (short and long loop DNCs) were confirmed by running agarose gel (0.8%) electrophoresis.

### Binding of biotin-labeled ssDNA with streptavidin

Two different approaches were used to bind the biotin-labeled ssDNA handles to streptavidin, one for short-loop DNC constructs and one long-loop DNC constructs. First, as shown in supplementary fig. 1, short-loop DNC labeled with biotin and digoxigenin at its terminals (8 μL of 100 pM) was incubated with streptavidin-coated silica beads (0.5 µL of 1% (w/v), 3.0 µm diameter) on the rotor for 30 minutes. The beads were washed two times with buffer (1x PBS, pH 7.4) and incubated with a mixture of strong and weak handles (1μL of 1 μM each) for 30 minutes. The beads were then washed with buffer three times to remove unbound DNA handles. Next, the washed beads were incubated with streptavidin (0.1 mg/mL) for 30 minutes and then excess of streptavidin was washed two times with buffer before the beads were injected into the chamber for measurements. Second, as shown in supplementary fig. 4, long-loop DNC labeled with digoxigenin at one end and with a 60 base long overhang on the other end was used for hairpin-barcoded measurements. The DNC was incubated with antidigoxigenin coated silica beads (0.5 µL of 1% (w/v), 3.0 µm diameter) on a rotor for 1 hour and washed two times before incubating with the strong handle and “no hairpin” weak handle (1μL of 0.2 μM each) for 30 minutes. The beads were washed with buffer three times and then incubated with streptavidin (0.1 mg/mL) for 30 minutes. Excess streptavidin was washed out by washing the beads two times with buffer and then then the beads were incubated with an equimolar mixture of short-hairpin and long-hairpin barcoded weak handles for 30 minutes. The excess DNA handles were washed out before injecting the beads into the chamber to carry out experiments.

### Measurement of distances between biotin-binding sites in streptavidin using DNC

The single-molecule mechanical experiments were performed in a home-built dual trap optical tweezers system^24,36,37^. Briefly, two optical traps were generated by splitting the 1064 nm laser beam into two using a polarizing beam splitter as described by Moffit et al.^38^. While one beam remained stationary, the second beam was controlled by a precisely positionable steerable mirror (Mad City Labs, Madison WI). The experiments with no barcodes were performed in phosphate buffered saline (1x PBS buffer, pH 7.4) supplemented with 0.01% (w/v) Roche Blocking solution (Roche) at room temperature (23 °C), while for barcoded distance measurements, the experimental buffer consisted of 1x PBS (pH 7.4) supplemented with 5 mM MgCl_2_ and 0.01% Roche block solution.

The sample chamber was assembled as discussed previously^24^. Briefly, double-sided Kapton tape (1 mm, DuPont) was sandwiched between a glass slide and a cover glass. Both the glass slides and cover glass were washed with 70% ethanol and dried with Argon flow. One side of the Kapton tape was attached to the surface of the glass slide and two channels leading to a reservoir with appropriate width were created by cutting the tape with a razor blade. Then the channel was covered by an argon-dried cover glass. Each channel and the reservoir were passivated with Roche Blocking solution (1%) for ∼30 min and then flushed with the experimental buffer before injecting the samples. For the “one distance measurement per streptavidin complex” experiment, a 1 µL aliquot of the above prepared DNC on the surface of streptavidin-coated beads was injected through one channel and 1 µL of the 0.1% (w/v) anti-digoxigenin coated silica beads (3 µm) were injected through the other channel. The openings were sealed with grease. The two different beads were separately trapped in the laser foci and one bead was moved closer to the other by using a steerable mirror to tether the DNC construct between the two optically trapped beads as shown in Fig. 2b. The tethered DNC was confirmed to be a single molecule by estimating the contour length of the whole caliper (∼1 µm) and by a single breakage of the tether. Calipers were confirmed to consist of a target streptavidin complex by observing a sudden change-in-extension due to shearing off of the weak handle.

After confirmation of a target molecule, the experiment was switched to force-jump mode between ∼0 pN and ∼26 pN, where the lower force allowed the rebinding of a weak handle and the higher force caused shearing of the weak handle. By repetitive cycling of the forces, large statistics of transitions were recorded for every individual caliper. The data were recorded at 1400 Hz using a custom Labview program (National Instruments Corporation, Austin, TX).

For experiments using barcoded handles to enable multiple distances to be measured on each single molecule complex, 1 µL of the above prepared beads with long-loop DNC was injected into one channel. The other beads were prepared by incubating streptavidin coated beads with biotin-labeled oligo that is complementary to the 60 base long overhang of DNC and 1 µL of these beads were injected into the other channel. The tether formation strategy was the same as previously mentioned except it was through hybridization of complementary sequences. A slightly different force-jump protocol compared to the previous measurement was used to enable reading of the barcodes from the force-extension traces. We first applied a slow ramping of the force up to ∼15 pN to probe the unfolding of a hairpin barcode, followed by a jump to a larger force (∼20 pN) to measure the distances of biotin-binding sites before jumping to an even higher force (∼30 pN) to shear-open the loop of DNC by releasing the weak handle, followed by a jump back to the distance measuring force (∼20 pN) (Fig. 3). This measurement cycle was repeated multiple times for each individual streptavidin-biotin complex to enable the collection of sufficient statistics for data analysis.

### Data analysis

Custom Labview software and custom-written MATLAB scripts were used to acquire the data and analyze it, respectively. IGOR Pro was used for statistical analysis and graphing of the measured data points. For the “one distance per complex” measurements, DNC unlooping was determined by the sudden change-in-length (*ΔL*) observed during the waiting time at the shearing force (Fig. 2 d and e). The compliance in *ΔL* due to the drop-in force caused by sudden unlooping was corrected by taking the slope tangent to the shearing force in the force *vs*. extension curve. In order to convert these changes in length (*ΔL*) to absolute distance measurements (*d*) we subtracted them from the effective loop length (*L*_*o*_) at the corresponding experimental conditions, i.e. *d* = *L*_*o*_ - *ΔL*. When making multiple distance measurements per streptavidin-biotin complex using hairpin barcodes, the length of the barcode signature was determined by measuring the change-in-extension due to unfolding of the hairpin. The section of the force vs extension curve before and after the unfolding transition was fitted with the Worm-like chain model and the change in contour length was determined. We measured the change-in-extension (*ΔL*) due to the unlooping transition of the DNC at ∼20 pN force by subtracting the length of caliper before and after unlooping at the same force. Then, the absolute distance between biotin-binding sites was determined by subtracting ΔL from the effective length of long-loop DNC calibrated at ∼20 pN force (Supplementary Figure 5) i.e. *d* = *L*_*o*_ - *ΔL*. The distances measured by unlooping are separated based on the clearly resolved barcodes (Supplementary Fig. 6).

## Author contributions

W.W. and W.S. conceived of the project. P.S., D.Y., W.W. and W.S. designed the experiments. P.S. conducted experiments, and P.S. and D.Y. performed data analysis. All authors discussed the results and analysis, and contributed to the manuscript, with the initial draft written by P.S. and W.W.

## Additional information

Supplementary information is available in the online version of the paper. Correspondence and requests for materials should be addressed to W.W.

## Data and materials availability

All relevant data are available from the supplementary files and the corresponding authors upon reasonable request.

