## Supplementary Materials for "Mapping Single-molecule Protein Complexes in 3D with DNA Nanoswitch Calipers"

**Supplementary Table 1**

List of DNA oligonucleotides that are used to synthesize DNA Nanoswitch Calipers (DNC), target DNA oligonucleotides and DNA handles used to label the peptides.

| Oligonucleotides | Sequences |
|---|---|
| BsaI-forward primer | AAA AAA GGT CTC GGC ATG ATA GCG CCC GGA AGA GAG |
| Biotin-reverse primer | /52-Bio/GGG TTC GTG CAC ACA GCC CAG CTT |
| Loop_short | ATG CCG CGT AAC CGG TCC GAG ACT CTC ACT GTG AAC TGA GTT TTT TTT TTT TTT TTT TTT TTT TTT TTT TTT TTT TTT TTT TTT TTT TTT TTT TTT TTT TTT TTT TTT TTT TTT TTT TCA GCG TCC GCG TGT CAG TGT |
| Loop_long | ATG CCG CGT AAC CGG TCC GAG ACT CTC ACT GTG AAC TGA GTT TTT TTT TTT TTT TTT TTT TTT TTT TTT TTT TTT TTT TTT TTT TTT TTT TTT TTT TTT TTT TTT TTT TTT TTT TTT TTT TTT TTT TTT TTT TTT TTT TTT TTT TTT TTT TTT TTT TTT TTT TTT TTT TTT TTT TTT TTT TTT TTT TTT CAG CGT CCG CGT GTC AGT GT |
| DNC1 | /5DigN/CGT CCC CAA ATC CGT ACC GGA TGC ACT GGC ACT GTG GGT GTG CGA CTT AAT TCC ATC CTG TTC CTG AGA CAA TAC AGC ACG ACC GCT TTC |
| DNC2 | /5Phos/TCT AAC TTA CAG AGC ATG GCG AAA GCG GTC GTG CTG TAT TGT CTC AGG AAC AGG ATG GAA TTA AGT CGC ACA CCC ACA GTG CCA GTG CAT CCG GTA CGG ATT TGG GGA CG |
| DNC3 | TCT CGG ACC GGT TAC GCG |
| Splint | AAG TTA GAA CAC TGA C |
| Surface DNA | Biotin - CTG AGT AAT GTG TAG GTA AAG ATT CAA AAG GGT GAG AAA GGC CGG AGA CAG TCA AAT CAC |
| DNC4 | GAT GGC CCG CTG AAA GGG CAG TGT TTC CCA GCG CCC TTC CTG GTA TGC GGA TTC TTT CGG GAG ATA GTA ATT AGC ACC TCC ATC GTC AGA CGG CGT CTG TGG CTC CGG CCT GAA CAG TGA |
| DNC5 | /5Phos//TCT AAC TTA CAG AGC ATG GCT CAC TGT TCA GGC CGG AGC CAC AGA CGC CGT CTG ACG ATG GAG GTG CTA ATT ACT ATC TCC CGA AAG AAT CCG CAT ACC AGG AAG GGC GCT GGG AAA CAC TGC CCT TTC AGC GGG CCA TCG TGA TTT GAC TGT CTC CGG CCT TTC TCA CCC TTT TGA ATC TTT ACC TAC ACA TTA CTC AG |
| GH-biotin | GCC ATG CTC TGT AAG TTA GAA CAC TGA CAC GCG GAC GCT GTT /3Bio/ |
| SH-biotin | /5Biosg/TTC TCA GTT CAC AGT G |
| SH-NoHP-biotin | /5Biosg/TTG TAT CAC TCC AAT GCT TTT TTT TTT TTT TTT TTT TTT TTT TTT TTT TTT TTT TTT TCT CAG TTC ACA GTG |

| | |
|---|---|
| SH-ShortHP-biotin | /5Biosg/TTT TTT TTT TTT TTT TTT TTT TTT TTT TCA TCC AAT GCT TTT TTT TGC ATT GGA TGT TCT CAG TTC ACA GTG |
| SH-LongHP-biotin | /5Biosg/TTT TTT TTT TTT TTG CTC CGA CGC TTT TTT TTT TTT TTT TTT TTT TGC GTC GGA GCT TCT CAG TTC ACA GTG |

Sequences from left to right are in the 5' to 3' direction.

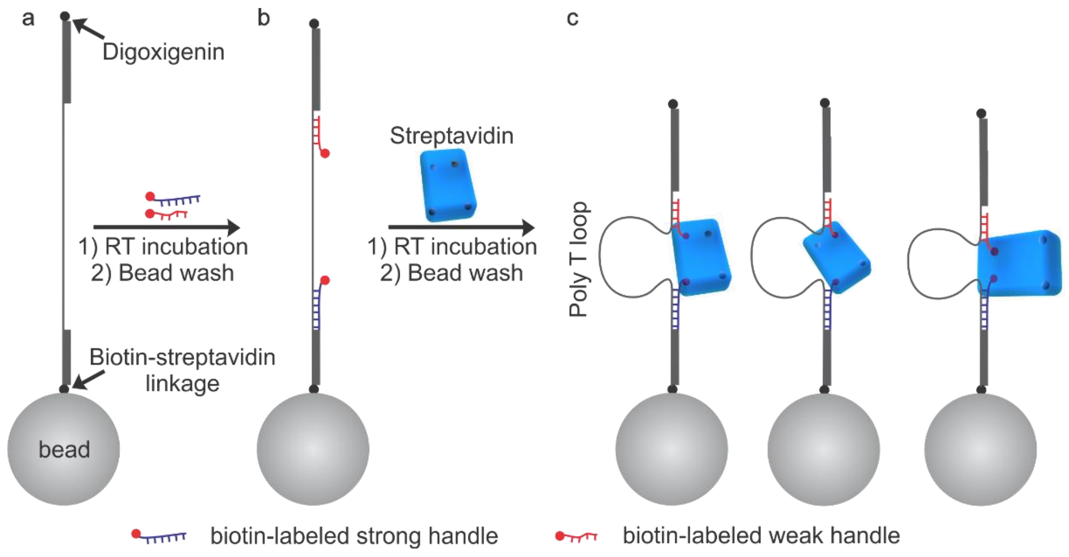


**Supplementary Figure 1.** Schematic outline of the synthesis of ssDNA labeled streptavidin for the measurement of one distance per streptavidin molecule. The distances are measured between two biotin-binding sites in each streptavidin molecule using DNA nanoswitch calipers in optical tweezers. **a,** DNA nanoswitch caliper is immobilized on the surface of streptavidin coated silica beads via biotin-streptavidin linkage. **b,** Washed from 'a' is incubated with a biotin-labeled long and strong DNA handle (purple) and a biotin-labeled short and weak DNA handle (red) to bind at their complementary sequences in DNC. **c,** Washed from 'b' is incubated with streptavidin to bind with the biotin ends of the DNA handles in DNC. Shown three constructs indicate the binding of two handles randomly into three different orientations.

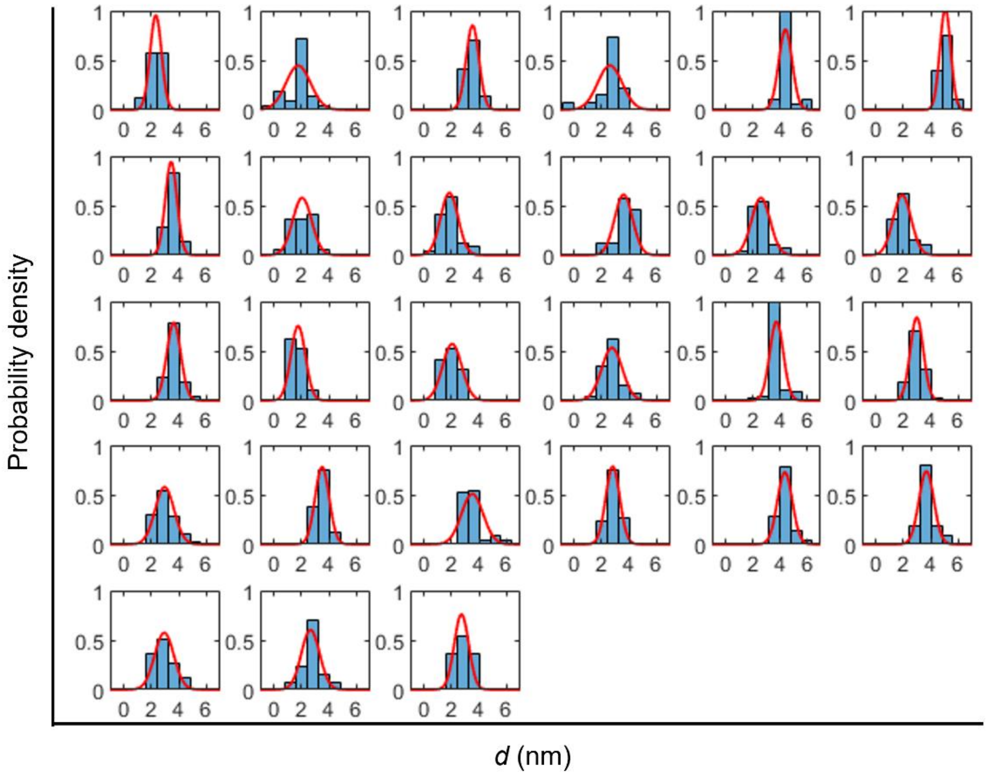


**Supplementary figure 2.** Histograms of the distances measured between two handles in every individual streptavidin molecule.

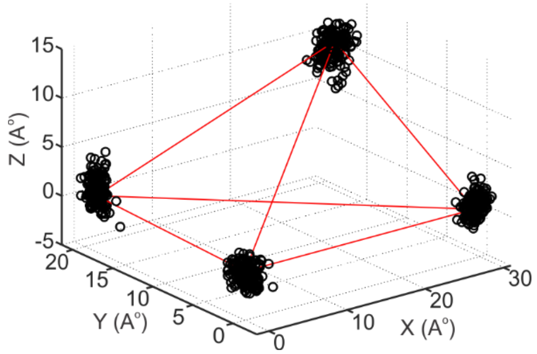


**Supplementary figure 3.** Reconstructed tetrahedron geometry (black circles) compared with the geometry of carbon in carboxylic group of biotins bound in streptavidin obtained by X-ray crystallography (red lines).

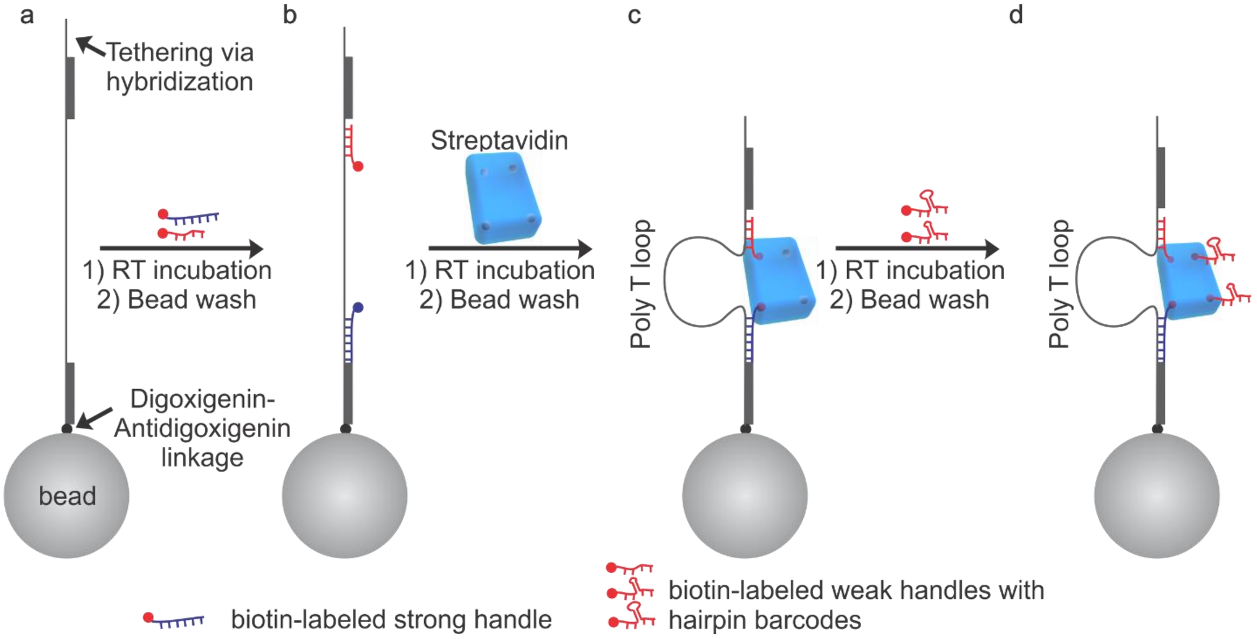


**Supplementary Figure 4.** Schematic outline of the synthesis of ssDNA labeled streptavidin for the measurement of all the three distances between biotin-binding sites using DNA nanoswitch calipers in optical tweezers. **a,** DNA nanoswitch caliper is functionalized on the surface of streptavidin coated silica beads via digoxigenin-antidigoxigenin linkage. **b,** Washed from 'a' is incubated with a biotin-labeled long and strong DNA handle (blue) and a biotin-labeled short and weak DNA handle (red) to bind at their complementary sequences in DNC. **c,** Washed from 'b' is incubated with streptavidin to bind with the biotin ends of the DNA handles. d, Washed from 'c' is incubated with two hairpin barcoded ssDNA handles.

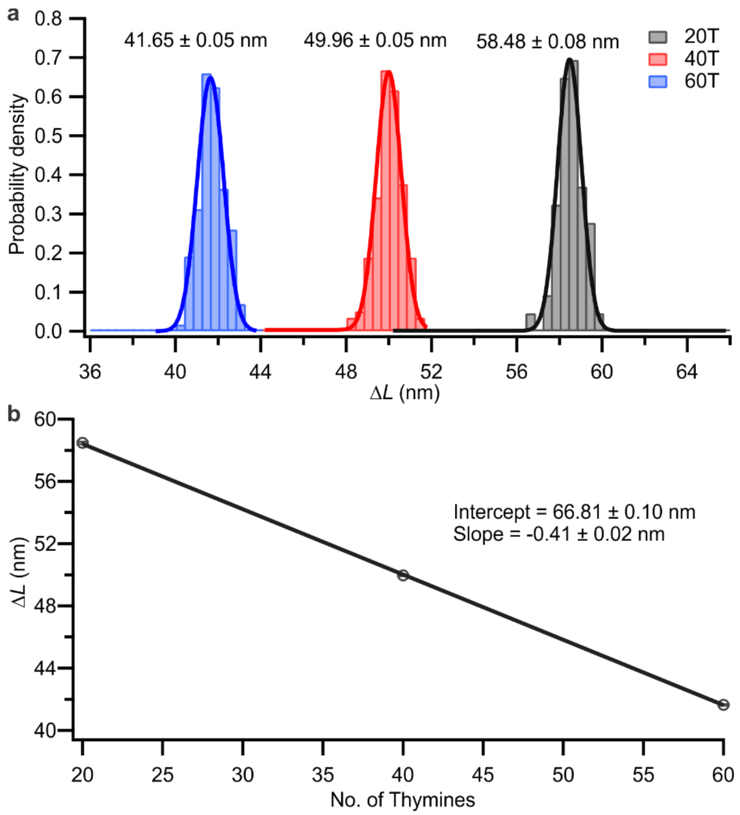


**Supplementary Figure 5. Calibration of long-loop DNA nanoswitch caliper at 20 pN. a,** Histograms of the change-in-extension measured for three different ssDNA target lengths; 20T, 40T and 60T. **b,** Correlation plot of the change-in-extension (*ΔL*) vs the length of ssDNA targets. Solid black line depicts the linear fit with $R^2 = 0.9946$. Error bars represent the calculated standard error of the mean.

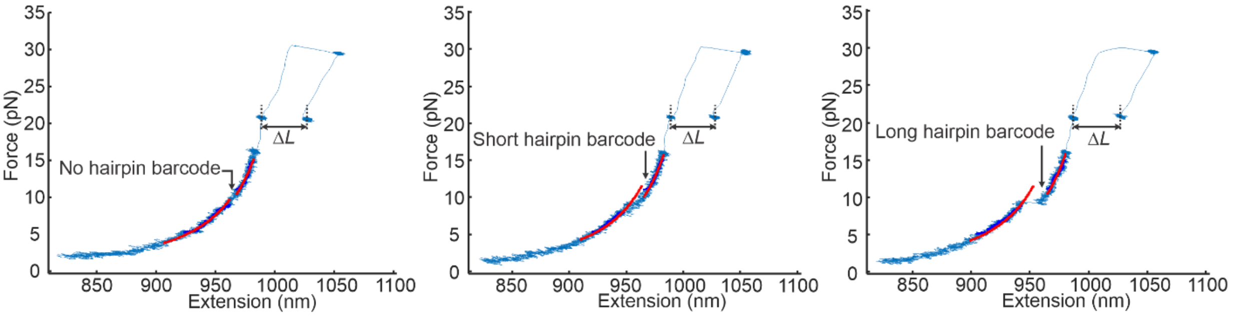


**Supplementary Figure 6.** Distance measurements in each cycle of measurement. Graphs from the Left to right panel are representative Force vs Extension curves in repeated cycles of measurement indicate the binding of all three barcodes in a single streptavidin molecule. A section of FX curve (dark blue) before and after unfolding of hairpin barcode is fitted with worm-like chain model[1] (solid red line) to measure the length of each barcode. The extension at ~20 pN force before and after unlooping of DNC determines the biotin-binding sites in streptavidin.

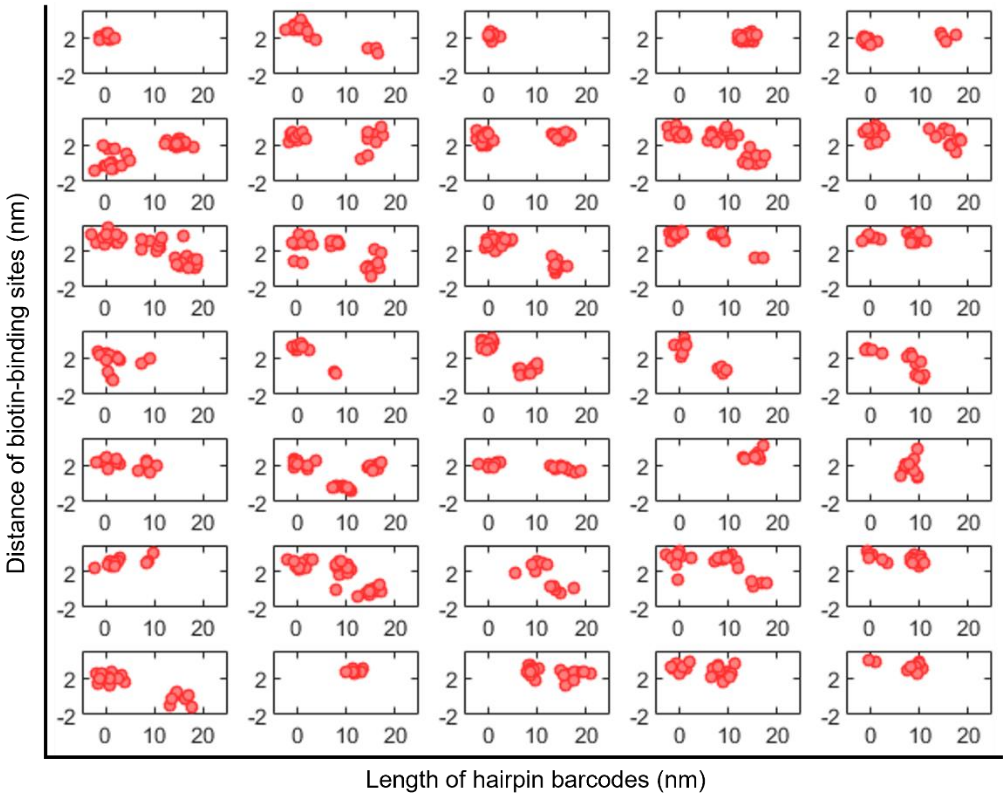


**Supplementary Figure 7.** Scatter plots of the measured length of hairpin barcodes and distance between the biotin-binding sites. Each plot represents the data acquired from a single molecule.

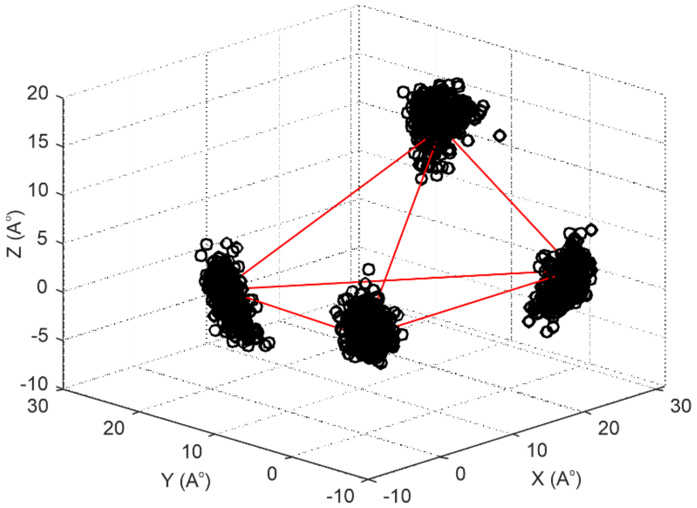


**Supplementary Figure 8.** Reconstructed tetrahedron geometry (black circles) from the measured distances (Figure 3g) compared with the geometry of carbon in carboxylic group of biotins bound in streptavidin obtained by X-ray crystallography (red lines).

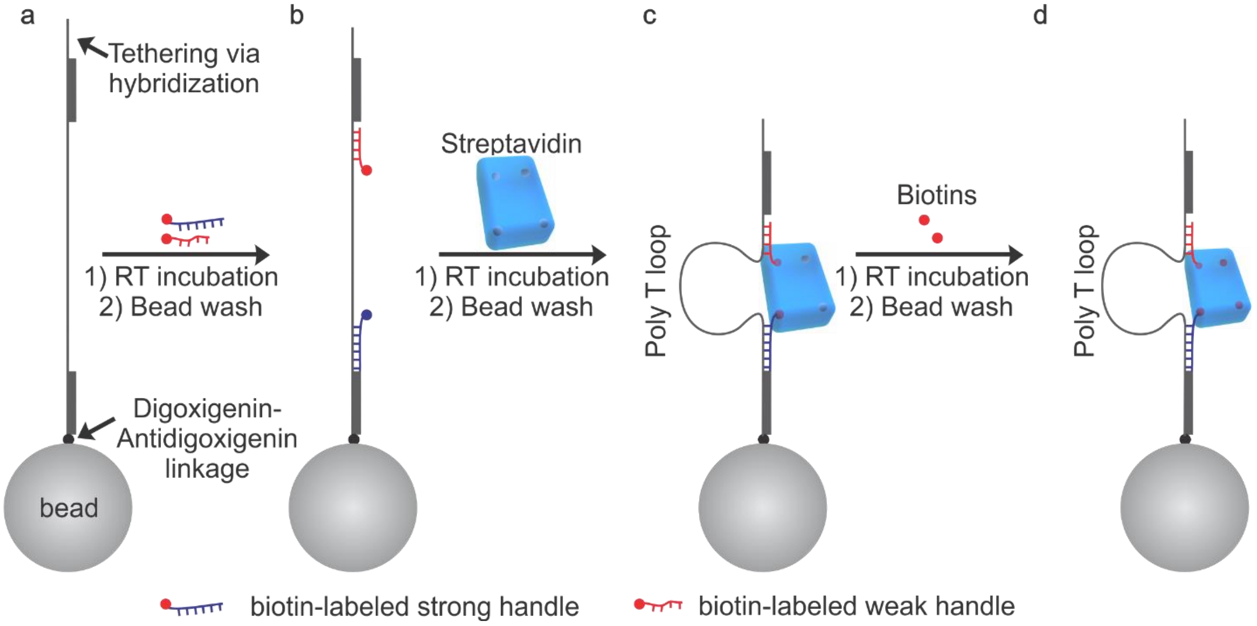


**Supplementary Figure 9.** Schematic outline of the synthesis of ssDNA labeled streptavidin for the measurement of one distance per streptavidin molecule that is fully occupied with biotins. **a,** DNA nanoswitch caliper is functionalized on the surface of streptavidin coated silica beads via digoxigenin-antidigoxigenin linkage. **b,** Washed from 'a' is incubated with a biotin-labeled long and strong DNA handle (purple) and a biotin-labeled short and weak DNA handle (red) to bind at their complementary sequences in DNC. **c,** Washed from 'b' is incubated with streptavidin to bind with the biotin ends of the DNA handles. **d,** Washed from 'c' is incubated with biotins to make fully occupied biotin-streptavidin complex.

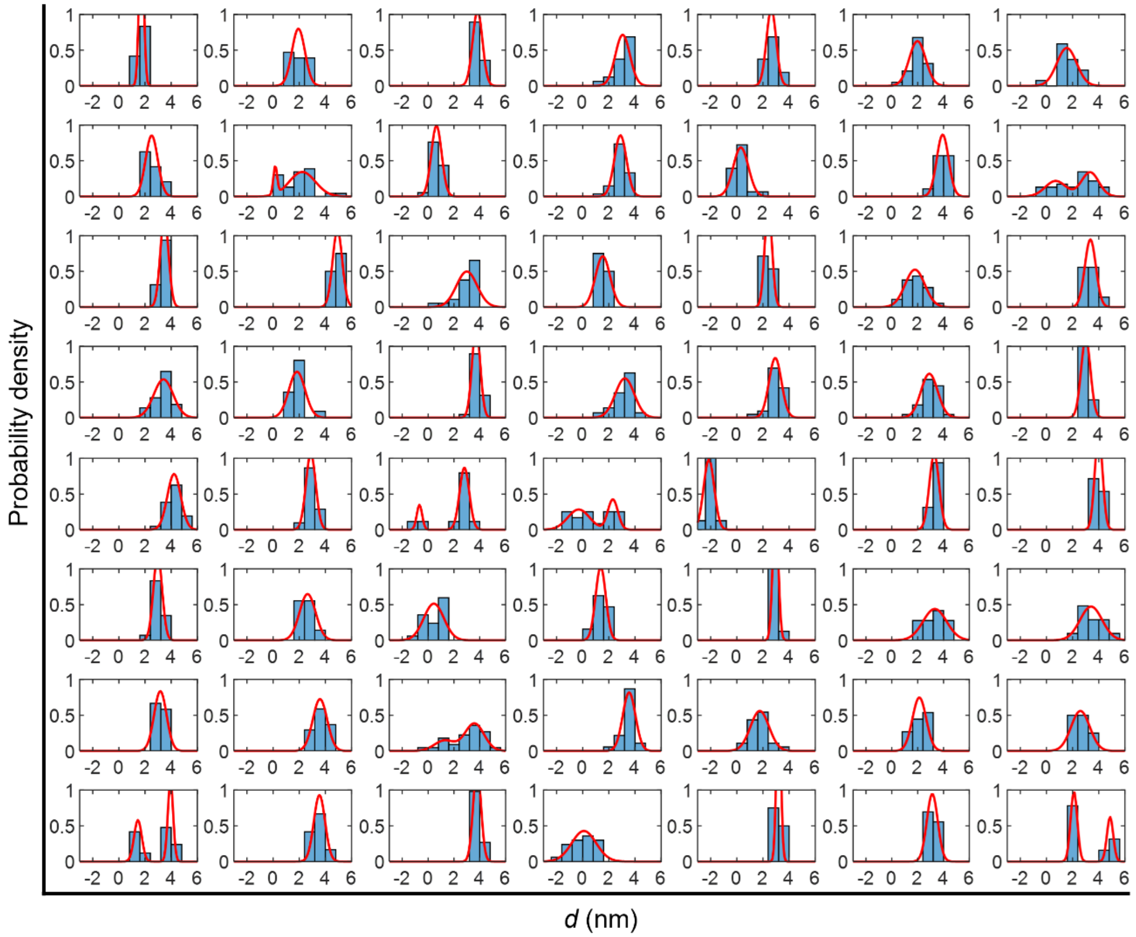


**Supplementary Figure 10.** Histograms of the distances measured between two handles in every individual streptavidin molecule that has the free-biotins bound into the rest of the two biotin-binding sites.

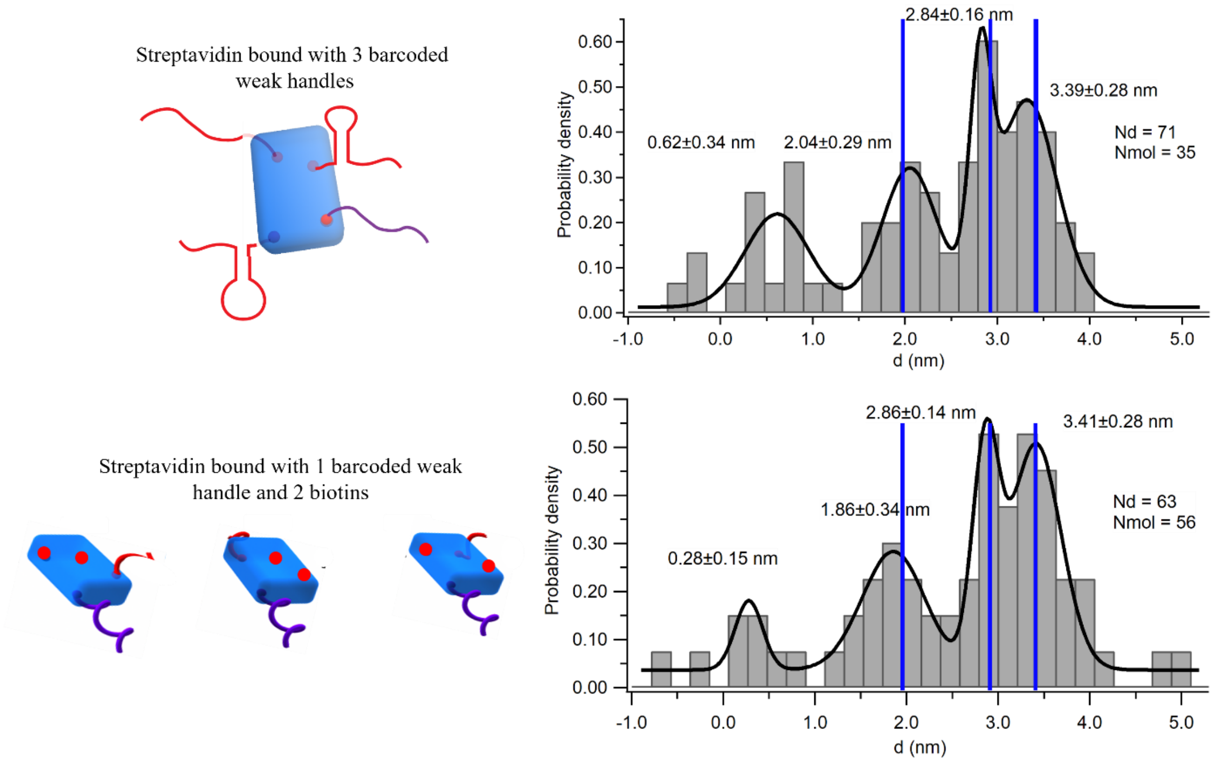


**Supplementary Figure 11.** Comparison of the measured distances between biotin-binding sites in streptavidin when it is fully bound with all the ssDNA handles (top panel) and two ssDNA handles and two biotins (bottom panel). Blue vertical lines depict the distances determined from the crystal structure.

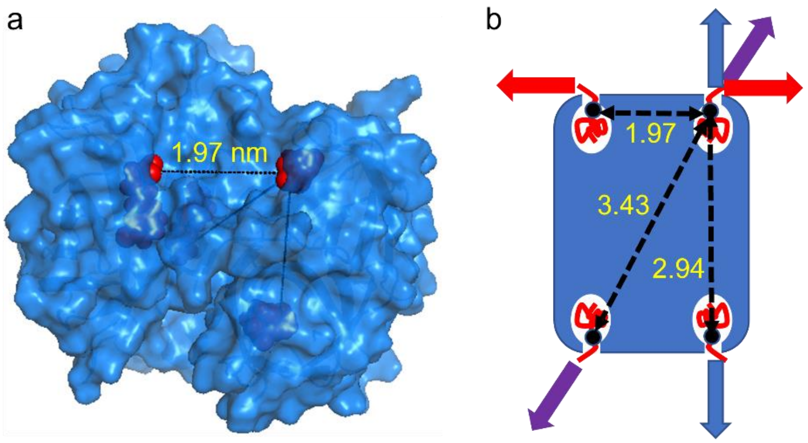


**Supplementary Figure 12. a,** Crystal structure of biotin-streptavidin complex indicating the slightly buried carboxylic group of biotins. **b,** The slightly buried carboxylic group of biotins have differential accessibility of measurements using DNC. The top right binding site is assumed to be binding with a strong handle and other binding sites have barcoded weak handles. Arrows indicate the direction of pulling and color codes are used to indicate the distance measurement across a specific pair of binding sites. The numbers depict the distances (nm) between the corresponding binding sites indicated by dotted lines.

**Supplementary Note 1.**

**Reconstruction of tetrahedron geometry from the measured distances**

X-ray crystallography and cryo-EM techniques have shown that the structure of the biotin-streptavidin complex has three-fold symmetries along all three axes[2,3] and the pairwise distances of the carboxylic acid groups in biotin can have only three unique values as shown in the following schematic figure.

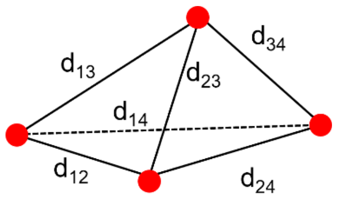


$$d_{12} = d_{34},\ d_{13} = d_{24}, \text{ and } d_{14} = d_{23}$$

By using DNC, we have measured these three distances in individual biotin-streptavidin complexes. To reconstruct the shape of tetrahedron using these distances, we first calculated the coordinates of the four corners using the following simple algebraic equations.

$$d_{12}^2 = (x_1 - x_2)^2 + (y_1 - y_2)^2 + (z_1 - z_2)^2$$
$$d_{13}^2 = (x_1 - x_3)^2 + (y_1 - y_3)^2 + (z_1 - z_3)^2$$
$$d_{14}^2 = (x_1 - x_4)^2 + (y_1 - y_4)^2 + (z_1 - z_4)^2$$

$x_1$, $y_1$, and $z_1$ are set to zero given the translational degree of freedom and rotational degree of freedom allows us to set one edge of the tetrahedron to be on the x-axis and another edge in the first quadrant of the xy-plane using the following equations.

$$x_2 = d_{12},\ y_2 = 0, \text{ and } z_2 = 0$$

$$z_3 = 0,\ x_3 > 0, \text{ and } y_3 > 0$$

This satisfies the condition that the overall volume of tetrahedron should be positive and produces two geometries that are mirror symmetries.
